# Developing a Petri dish test to detect resistance to key herbicides in *Lolium rigidum*

**DOI:** 10.1101/2021.04.08.438914

**Authors:** Martina Badano Perez, Hugh J Beckie, Gregory R Cawthray, Danica E Goggin, Roberto Busi

**Author notes:** Author for correspondence: Roberto Busi, Australian Herbicide Resistance Initiative, School of Agriculture and Environment, University of Western Australia, Perth, Australia.

## Abstract

Overreliance on herbicides for weed control is conducive to the evolution of herbicide resistance. Annual ryegrass (*Lolium rigidum* Gaud.) is a species that is prone to evolve resistance to a wide range of herbicide modes of action. Rapid detection of herbicide-resistant weed populations in the field can aid farmers to optimize the use of herbicides for their control. The feasibility of a portable agar-based test to rapidly and reliably detect annual ryegrass resistance to key herbicides such as clethodim, glyphosate, pyroxasulfone and trifluralin on-farm was investigated. The three research phases of this study show that: a) easy-to-interpret results are obtained with non-dormant seed from well-characterised susceptible and resistant populations, and resistance is detected as effectively as with traditional dose-response pot-based resistance assays. However, the test may not be suitable for portable use on-farm because of b) the low stability of some herbicides such as trifluralin and clethodim in agar and c) the tendency of seed dormancy in freshly-harvested seeds to confound the results. The agar-based test is best used as a research tool as a complement to confirm results obtained in traditional pot-based resistance screenings. Comprehensive agar test and / or whole-plant resistance tests by herbicide application at the recommended label rate (whole plants grown in pots) are the current benchmark for proactive in- and off-season resistance testing and should be promoted more widely to allow early detection of resistance, optimization of herbicide technology use and deploy appropriate weed management interventions.

## 1 Introduction

From the 1880s, highly adaptable annual or rigid ryegrass (*Lolium rigidum* Gaud.) populations were introduced to different environments in southern Australia to use as a pasture for livestock grazing (Kloot, 1983). After a substantial move towards cropping in the 1950s, the control of annual ryegrass infestations became a priority in wheat, barley and canola because of the highly competitive and economically-damaging nature of the weed even at the early developmental stage of two leaves (Smith and Levick, 1974). Weeds such as annual ryegrass are a major constraint for Australian farming systems; the financial cost from yield loss and weed control expenditure accounts for AU $3.3 billion per year (Llewellyn et al., 2016). Effective control of ryegrass relies on systematic herbicide use as a key component of crop protection in the Australian minimal-tillage dryland farming systems, enabling producers to sow crops in fields with stubble retention and avoid erosive and labour-intensive soil cultivation (D’Emden and Llewellyn, 2006). However, this high reliance on herbicides to control weeds promotes the evolution of herbicide-resistant genotypes (Kreiner et al., 2018). The most challenging species is annual ryegrass, due to (1) its propensity to rapidly evolve resistance to a wide range of herbicides with different modes of action (Broster et al., 2019; Heap, 2020; Owen et al., 2014b), and (2) the highly variable dormancy level of its seeds, allowing different cohorts to germinate across the growing season and avoid knockdown (burndown) and/or in-crop herbicide application (Somerville et al., 2017).

Generally, the discovery of resistant weed populations occurs in the field as a consequence of the failure of a herbicide treatment that was previously effective. If resistance could be detected at an early stage, by proactively scouting fields and screening weed populations routinely, it may be possible to anticipate the causes of herbicide resistance and thereby propose changes in management designed to achieve effective weed control and minimize the spread of resistance (Beckie et al., 2000; Norsworthy et al., 2012).

Proactive resistance testing is not a commonly-adopted practice among farmers, however, because testing large numbers of samples for resistance using traditional pot-based methods is expensive, time-consuming, and requires a large amount of space (Kaundun et al., 2011). To encourage farmers to test for resistance more frequently, it is necessary to provide them with step-by-step guidelines from seed collection to seed testing (Burgos, 2015). Even more important is to develop a faster and less expensive diagnostic technique, with simple interpretation of results. Methods involving germination of seeds or pollen on herbicide-containing media, incubation of leaf samples in herbicide solutions, spraying of separated grass weed tillers or measurement of hydroponically-grown seedling development in the presence of herbicide have been used in research laboratories (Burgos et al., 2013), but none have been developed for use by the farmers themselves to enable on-farm resistance testing.

The aim of this study was to develop and evaluate an easy-to-interpret on-farm test to rapidly detect resistance to both pre- and post-emergence herbicides in annual ryegrass populations by simply placing seeds on herbicide-containing agar and observed seed germination after a few days. The herbicide impregnated-agar would be prepared in a laboratory with a view to distributing testing kits to the end-user. For this reason we measured stability of the herbicides in the agar in order to assess the potential portability and shelf-life of the test. The significance of seed dormancy as a barrier to the success of the test (Beckie et al., 2000) was also assessed.

## 2 Materials and methods

### 2.1 Plant material

Well-characterized annual ryegrass populations, with little to no dormancy (>96% germination within 7 d), were used to optimize the agar-based resistance test and its correlation to traditional whole-plant pot-based experiments (Table 1). These populations have been characterized as susceptible or resistant in standard pot-based dose-response experiments using clethodim, glyphosate, pyroxasulfone and trifluralin. An additional 100 populations collected from different fields in Western Australia in 2018 and 2019 (Busi et al., 2020) were used to validate the correlation between an agar test and a whole-plant test in pots. Before testing, seed dormancy in these populations was relieved by hydrated storage in the dark at room temperature for 14 d. (Steadman, 2004) Populations selected for low (conditional) dormancy, requiring alternating light and temperature to trigger germination, or very high primary dormancy, requiring extensive after-ripening or dark-stratification to sensitise them to germination cues (Steadman, 2004), were also included in some experiments (Goggin et al., 2010) (Table 1).

**Table 1.**
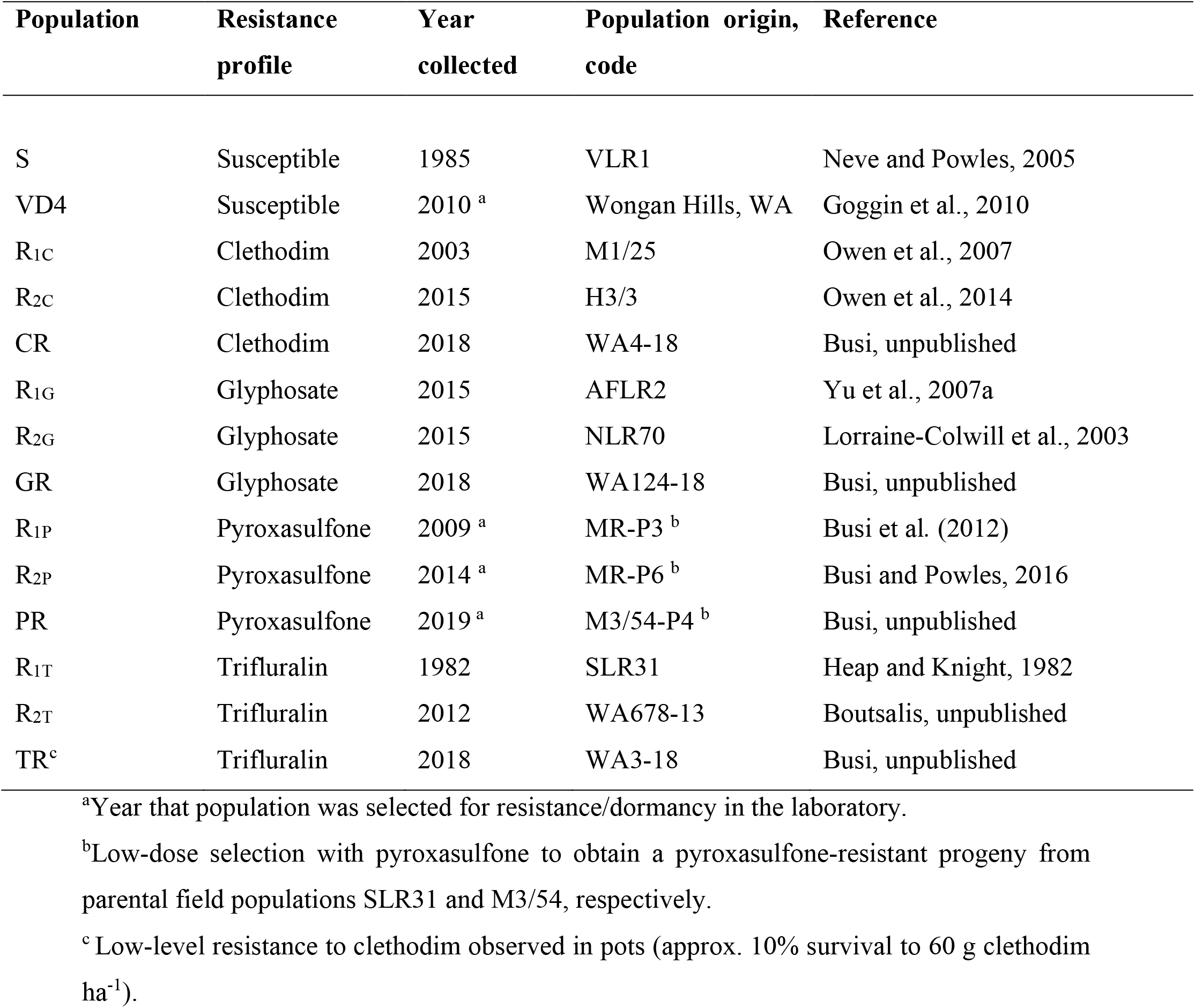
Populations of known resistance status used to optimize the agar test.

| Population | Resistance profile | Year collected | Population origin, code | Reference |
| --- | --- | --- | --- | --- |
| S | Susceptible | 1985 | VLR1 | Neve and Powles, 2005 |
| VD4 | Susceptible | 2010 <sup>a</sup> | Wongan Hills, WA | Goggin et al., 2010 |
| R<sub>1C</sub> | Clethodim | 2003 | M1/25 | Owen et al., 2007 |
| R<sub>2C</sub> | Clethodim | 2015 | H3/3 | Owen et al., 2014 |
| CR | Clethodim | 2018 | WA4-18 | Busi, unpublished |
| R<sub>1G</sub> | Glyphosate | 2015 | AFLR2 | Yu et al., 2007a |
| R<sub>2G</sub> | Glyphosate | 2015 | NLR70 | Lorraine-Colwill et al., 2003 |
| GR | Glyphosate | 2018 | WA124-18 | Busi, unpublished |
| R<sub>1P</sub> | Pyroxasulfone | 2009 <sup>a</sup> | MR-P3 <sup>b</sup> | Busi et al. (2012) |
| R<sub>2P</sub> | Pyroxasulfone | 2014 <sup>a</sup> | MR-P6 <sup>b</sup> | Busi and Powles, 2016 |
| PR | Pyroxasulfone | 2019 <sup>a</sup> | M3/54-P4 <sup>b</sup> | Busi, unpublished |
| R<sub>1T</sub> | Trifluralin | 1982 | SLR31 | Heap and Knight, 1982 |
| R<sub>2T</sub> | Trifluralin | 2012 | WA678-13 | Boutsalis, unpublished |
| TR<sup>c</sup> | Trifluralin | 2018 | WA3-18 | Busi, unpublished |

<sup>a</sup>Year that population was selected for resistance/dormancy in the laboratory.
<sup>b</sup>Low-dose selection with pyroxasulfone to obtain a pyroxasulfone-resistant progeny from parental field populations SLR31 and M3/54, respectively.
<sup>c</sup> Low-level resistance to clethodim observed in pots (approx. 10% survival to 60 g clethodim ha<sup>-1</sup>).

### 2.2 Optimization of the agar test

Commercial formulations of herbicide were added to liquid 0.6% (w/v) agar and poured to a depth of 1 cm into 5-cm deep containers of 12.5 cm diameter. The herbicide concentrations chosen for dose-response experiments were based around discriminating doses identified in preliminary experiments and in the literature (Beckie et al., 2000; Chen et al., 2018b; Kaundun et al., 2011). Clethodim (Sequence, 240 g a.i. L^−1^; Nufarm Australia) was used at concentrations of 0.125, 0.25, 0.5, 1, 2, 3, 4 and 5 μM; glyphosate (Weedmaster ARGO, 540 g a.i. L^−1^; Nufarm Australia) at 10, 100, 200, 300, 400, 500, 600 and 1000 μM; pyroxasulfone (Sakura, 850 g a.i. kg^−1^; Bayer Crop Science) at 37.5, 75, 150, 300, 600 and 1200 nM; and trifluralin (Triflur X, 480 g a.i. L^−1^; Nufarm Australia) at 5, 10, 20, 25, 30, 50 and 100 μM. Herbicide-free controls were included in every experiment. Twenty five seeds per population per treatment were placed on the surface of the herbicide-impregnated agar and incubated in a growth chamber at 25/15°C day/night with a 12 h photoperiod of cool white fluorescent light (30 – 60 μmol m^−2^ s^−1^) (Goggin et al., 2010) for 7 d. Individual seedlings were considered resistant if they could grow actively, with the length of their coleoptiles reaching ≥ 4 cm. The agar container was the experimental unit. Three replicates of each herbicide concentration were prepared and the experiment was repeated twice at the University of Western Australia. The minimum dose which killed 100% of the susceptible individuals was considered to be the discriminating dose.

### 2.3 Pot-based dose-response experiments

The results of the agar test were compared with those of a traditional pot-based resistance test by conducting dose-response experiments in 6 × 6 × 6 cm plastic cells filled with potting mix (25% peat moss, 25% washed river sand and 50% mulched pine bark). For the pre-emergence herbicides (pyroxasulfone and trifluralin), 50 seeds per cell were sprayed while situated on the soil surface and then covered with a thin layer of fresh potting mix. For the post-emergence herbicides (clethodim and glyphosate), 50 seedlings in each cell were sprayed at the two-to three-leaf stage. Herbicides were applied using a cabinet track sprayer with flat-fan nozzles, delivering 216 L ha^−1^ per pass at 200 kPa of pressure (Owen et al., 2014b). The range of doses applied was based around the recommended field rate for each herbicide (in g a.i. ha^−1^: clethodim, 120; glyphosate, 540; pyroxasulfone, 100; trifluralin, 960) or on previous studies published (Busi et al., 2012; McAlister et al., 1995; Owen et al., 2014b; Powles et al., 1998; Yu et al., 2007b). Clethodim was applied at 62, 125, 250, 500 and 750 g ha^−1^; glyphosate at 270, 540, 810, 1080 and 1350 g ha^−1^; pyroxasulfone at 12, 25, 50, 100, 150 and 200 g ha^−1^; and trifluralin at 60, 120, 240, 480 and 720 g ha^−1^. Unsprayed controls were also included. Plants were grown in the glasshouse in autumn (March/April) with regular watering and fertilizer application, and survival was assessed at 21 d after the spray application. Similarly to the agar test, survivors were those plants that could establish, develop two new leaves and grow actively. Each cell was the experimental unit. There were three replicates of each treatment, and each experiment was performed twice.

### 2.4 Validation of the agar test with uncharacterized populations

One hundred uncharacterized populations collected from the field in 2018/19 were incubated on agar containing the discriminating herbicide doses identified in the optimization phase of the experiment. To account for any seed dormancy remaining after 14-d dark stratification, sub-samples were germinated on control agar lacking herbicide. The seed germination and survival of treated seeds were expressed as percentage of the untreated control. Twenty five seeds per population per treatment were used with one replicate per population. The populations were also sprayed with the recommended rate of each herbicide in pot-based tests to determine the correlation between observed resistance levels in agar and pots. Twenty five seeds per population per treatment were used with one replicate per population. Plant survival, classified as the percentage of seedlings that had produced at least two new leaves after herbicide exposure, was assessed at 21 d after treatment.

### 2.5 Investigation of interference from seed dormancy

To determine if dormant ryegrass seeds can be forced to germinate within the 7-d time frame of the agar test, the well-characterised low- and high-dormancy populations (ND4 and VD4) from Goggin et al. (2010) were subjected to a range of potential dormancy-breaking treatments (see Supplementary Table S1 for full details and references). Seeds were then incubated on agar under either controlled conditions (25/15°C, 12 h photoperiod) or next to the laboratory window to simulate conditions in a farmer’s house. Germination was counted every 7 d for 42 d, and dead or empty seeds were excluded from calculations. There were three replicates of each treatment.

The effect of dark stratification in the presence of herbicides was assessed using population S (Table 1) as a susceptible, non-dormant control; the very-dormant population (VD4) as a susceptible, dormant control; and freshly-harvested (not after-ripened) samples from five populations with known resistance to clethodim (population CR), glyphosate (GR), pyroxasulfone (PR) or trifluralin (TR) (Table 1). Seeds (10 per treatment per population) were imbibed on agar as previously described and placed under environmental conditions of 25/15°C in a growth cabinet or on the laboratory window sill for 42 d, or first dark-stratified for 21 d (dishes wrapped in foil and placed in a controlled 20°C room or on the laboratory window sill) before being transferred to germination conditions for a further 21 d. The agar contained herbicide during stratification and germination, or during germination only, as indicated in the experimental scheme presented in Figure 3A. Sub-lethal herbicide treatments were included because of the longer incubation time of the seeds with the herbicides. Germination was recorded every 7 d. For the purposes of this experiment, which assessed herbicide resistance as well as seed germination, a coleoptile length of ≥ 4 cm was required for a seed to be counted as germinated. There were three replicates of each treatment. The experiment was repeated once.

### 2.6 Herbicide stability in agar

Technical-grade herbicides were used for the initial optimisation of high-performance liquid chromatography (HPLC) methods. Stock solutions of 1 mM clethodim, pyroxasulfone or trifluralin in 50% acetonitrile were prepared and dilutions then made to 10 – 200 μM, 5 – 10 μL of which were directly injected onto a Nova-Pak C_18_ column (150 mm × 3.9 mm i.d.) with 4 μm particle size (Waters, Milford, MA, USA) attached to a 600E dual-head pump with 717 Plus autosampler and 996 photodiode array detector (Waters). Column temperature was held at 30°C and samples in the autosampler kept at 15°C. Chromatographic separation was achieved with a flow rate of 1 mL min^−1^ and a linear gradient of 40 – 90% (v/v) acetonitrile in 0.1% (v/v) formic acid over 5 min, then held for 0.5 min before immediate change to 100% acetonitrile, and subsequently held for 5 min before returning to the original conditions of 40% acetonitrile for 10 min prior to the next injection (Sandín-España et al., 2016). Absorbance at 255 nm was used for quantification of herbicides. Under these conditions, the retention times for each herbicide were: pyroxasulfone, 7.8 min; clethodim, 10.8 min; and trifluralin, 11.4 min. Positive identification of herbicides was made by comparing standard retention time and photodiode array (PDA) peak spectral analyses across 200 – 500 nm.

Glyphosate (1 mM in water) was derivatised with FMOC according to Ibáñez et al. (2005) and 10 μL injections were separated at a flow rate of 1 mL min^−1^ on the same C_18_ column as above. Column temperature was held at 35°C and samples in the autosampler kept at 15°C. The mobile phase was a gradient of ammonium acetate (pH 5.02) and acetonitrile adapted from Ibáñez et al. (2005), with initial conditions of 10% acetonitrile followed by immediate change to 35% acetonitrile after 0.1 min, then held at 35% for 10 min before immediate change back to 10% for column equilibration prior to the next injection. Derivatised glyphosate was detected with a 996 photodiode array detector (Waters) at 265 nm. Under these conditions, the retention time for glyphosate was 7.3 min. Positive identification of derivatised glyphosate was made by comparing standard retention time and PDA peak spectral analyses across 210 – 350 nm.

Subsequent HPLC analysis of the commercially formulated herbicides at the same concentration of active ingredient showed that there was no interference with the herbicide peaks from other compounds present in the formulations (data not shown). The detection limit was approximately 0.2 nmol per injection.

Stock solutions of each formulated herbicide were prepared by diluting in water to a concentration of 10 mM active ingredient. These were stored in glass bottles at 4°C in the dark and aliquots were removed at regular intervals and stored at −20°C for HPLC analysis of two technical replicates. The stability of each formulated herbicide when dissolved in a 0.6% (w/v) agar matrix was tested by preparing agar containing 2 μM clethodim, 400 μM glyphosate, 150 nM pyroxasulfone or 20 μM trifluralin. Dishes of agar were prepared for storage at (i) ambient laboratory temperature (mean, 21°C) and light (mean, 50 μmol m^−2^ s^−1^ during the day); (ii) ambient laboratory temperature in the dark (dishes wrapped in foil); or (iii) 4°C in the dark, with three replicates for each. Pilot studies on samples stored in the 25/15°C growth cabinet showed that there was no difference in herbicide degradation under ambient laboratory conditions compared to the growth cabinet (data not shown). The total mass of the agar in each dish was recorded, and then samples were taken using a 12 mm cork-borer at 0, 5 and 10 d after preparation. The mass of each sample was recorded and the agar plugs were then immediately chopped into 1 mm^3^ pieces and extracted by vigorous agitation in 100% acetonitrile (clethodim, pyroxasulfone and trifluralin) or in water (glyphosate). Samples were then centrifuged at 18,000 *g* for 10 min to pellet the agar. The supernatant volumes were recorded and the trifluralin and glyphosate samples were immediately stored at −80°C. The pyroxasulfone and clethodim samples were concentrated under a stream of air, partitioned against an equal volume of ethyl acetate to remove the water, evaporated to dryness and then redissolved in 30 – 50 μL of 100% acetonitrile and stored at −80°C until HPLC analysis was performed.

### 2.7 Statistical analysis

Dose-response curves were constructed for each population, herbicide and test system (agar versus pot test). Data from repeated experiments were pooled and subjected to non-linear regression analysis, using a log-logistic model with three parameters:

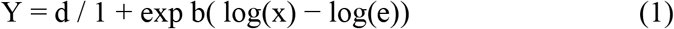

Y corrresponds to the survival response to herbicides for each population, d is the upper limit, b is the slope of the curve, x is the herbicide dose, and e is the dose causing 50% mortality response (Ritz and Streibig, 2005). Thus, the lethal dose required to kill 50% of the the population (LD_50_) and respective 95% confidence intervals were estimated for each herbicide and each population from dose-response analysis. The resistance index (RI) was calculated as the ratio of the LD_50_ of the resistant population and standard susceptible population S. Correlation analysis was conducted on plant survival percentage values obtained in 100 annual ryegrass populations, tested at the discriminating dose in agar tests versus the recommended herbicide dose in pots. Pearson’s correlation coefficient (r) and two-tailed P-values were estimated for pair wise combinations of plant survival observed in agar and pots for each of the four herbicides tested in the study. The statistical analysis was conducted with GraphPad Prism (GraphPad Software, Inc., La Jolla, California). Herbicide stability and seed germination data were analysed using one-factor ANOVA and Fisher’s LSD test with all assumptions held under square root arcsine transformations. Back-transformed data are presented in all figures.

## 3 Results and Discussion

The aim of this proof-of-concept study was to find a simplified and accurate alternative to traditional pot-based herbicide resistance screening experiments. An agar-based test was developed which could be prepared under carefully controlled conditions and then distributed to farmers, consultants or any other user to detect herbicide resistance in the field by simply placing weed seeds into a container and observing coleoptile elongation after a week. Annual ryegrass was identified as an ideal and suitable model weed species due to its reported capacity to evolve resistance to multiple herbicide modes of action (Heap, 2020) and its prevalence in Australian cropping systems.

### 3.1 Development of the agar test and comparison with the pot experiment

The dose-response experiment in agar with the four herbicides showed that it was possible to visually discriminate between susceptible and resistant annual ryegrass populations and individuals (Figure 1). The discriminating range of concentrations was identified for all four herbicides where they caused 100% mortality and full inhibition of coleoptile elongation in the S population but >1% survival and growth in the R populations. Clethodim at 0.25 – 1 μM resulted in 4 – 52% survival in the two clethodim-resistant populations (Figure 2A) and for glyphosate at 400 – 600 μM a survival response ranging between 6 and 59% was observed for the glyphosate-resistant populations (Figure 2B). For pyroxasulfone, the discrmininating concentrations (0.075 – 0.15 μM) resulted in 2 – 20% survival of the pyroxasulfone-resistant populations (Figure 2C), whilst survial to trifluralin was relatively high in both trifluralin-resistant populations, ranging between 10 and 60% (Figure 2D). The dose-response experiment in pots showed the expected discrimination between the susceptible and resistant populations, with the former being 100% killed at the recommended field rate of each herbicide (data not shown). It is clear from the statistical analysis of calculated LD_50_ values and interpretation of Resistance Indices that herbicide resistance can be similarly and effectively identified with either the agar or traditional whole plant screening method (Table 2).

**Table 2.** Lethal dose required to kill 50% (LD50) of each population in the agar test and pot experiment. LD50 values (with 95% confidence intervals in parentheses) were calculated from a three paramenter non-linear regression model. The resistance index (RI) was calculated based on the ratio of LD50 of the R populations to the susceptible population S. All estimated LD50 values of the R populations were significantly different from S (*P*<0.05).

| Herbicide | Population | Agar test (μM) |  | Pot study (g ha<sup>-1</sup>) |  |
| --- | --- | --- | --- | --- | --- |
|  |  | LD<sub>50</sub> (95% CI) | RI | LD<sub>50</sub> (95% CI) <sup>a</sup> | RI |
| Clethodim | R<sub>1C</sub> | 0.23 (0.13 - 0.43) | 4.6 | > 750 | > 17 |
| Clethodim | R<sub>2C</sub> | 0.18 (0.05 - 0.76) | 3.6 | > 750 | > 17 |
| Clethodim | S | 0.05 (0.004 to 0.007) | -- | 45 (21.4 – 68.5) | -- |
| Glyphosate | R<sub>1G</sub> | 370 (189 – 1110) | 4.0 | >1350 | > 7 |
| Glyphosate | R<sub>2G</sub> | 347 (164.3 - 770.2) | 3.8 | >1350 | > 7 |
| Glyphosate | S | 91.4 (37.5 - 183.5) | -- | 184 (143 – 225) | -- |
| Pyroxasulfone | R<sub>1P</sub> | 0.06 (0.03 - 0.095) | 15 | 46 (26.4 – 65.6) | 7.7 |
| Pyroxasulfone | R<sub>2P</sub> | 0.03 (0.015 - 0.06) | 7.5 | 22 (14.1 – 29.8) | 3.7 |
| Pyroxasulfone | S | 0.004 (0.0004 - 0.007) | -- | 6 (2.1 – 9.9) | -- |
| Trifluralin | R<sub>1T</sub> | 35 (6 – 391) | 29 | > 720 | > 8 |
| Trifluralin | R<sub>2T</sub> | 5.4 (0.3 – 41) | 4.5 | 203 (54 – 352) | 2.1 |
| Trifluralin | S | 1.2 (0.6 – 1.8) | -- | 96 (80.3 – 111.7) | -- |

<sup>a</sup> If an LD<sub>50</sub> value could not be calculated due to >50% survival at high doses, the LD<sub>50</sub> value was designated as greater than the highest herbicide dose tested.

**Figure 1.**
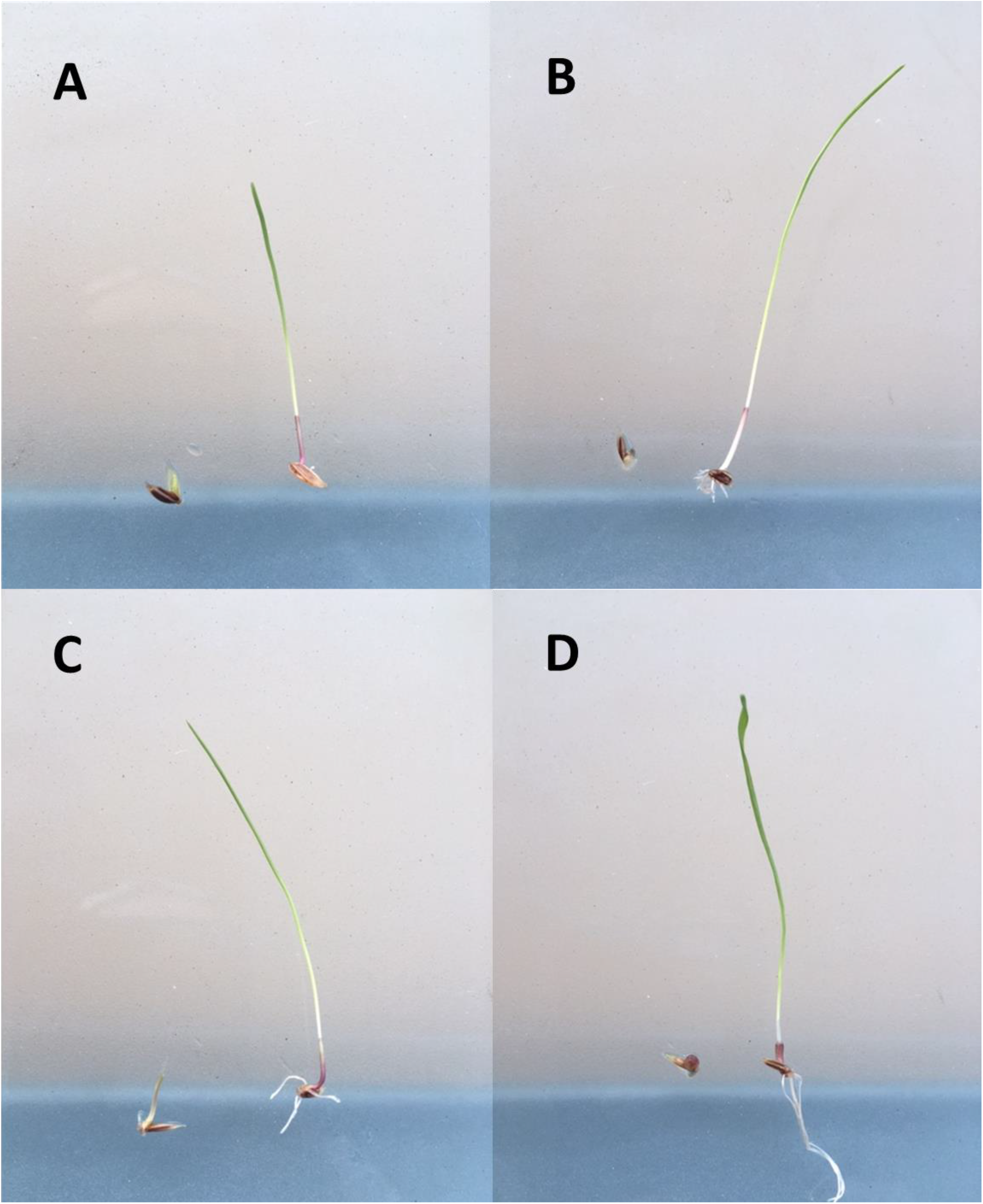
Inhibition of seed germination and development was used to discriminate herbicide susceptibility (left) from herbicide resistance (right) in annual ryegrass (*Lolium rigidum*). Herbicide resistance could be detected after 7 d with non-dormant seed by deploying a single herbicide concentration of A) clethodim [1 μM], B) glyphosate [400 μM], C) pyroxasulfone [0.15 μM] and trifluralin [25 μM].

**Figure 2.**
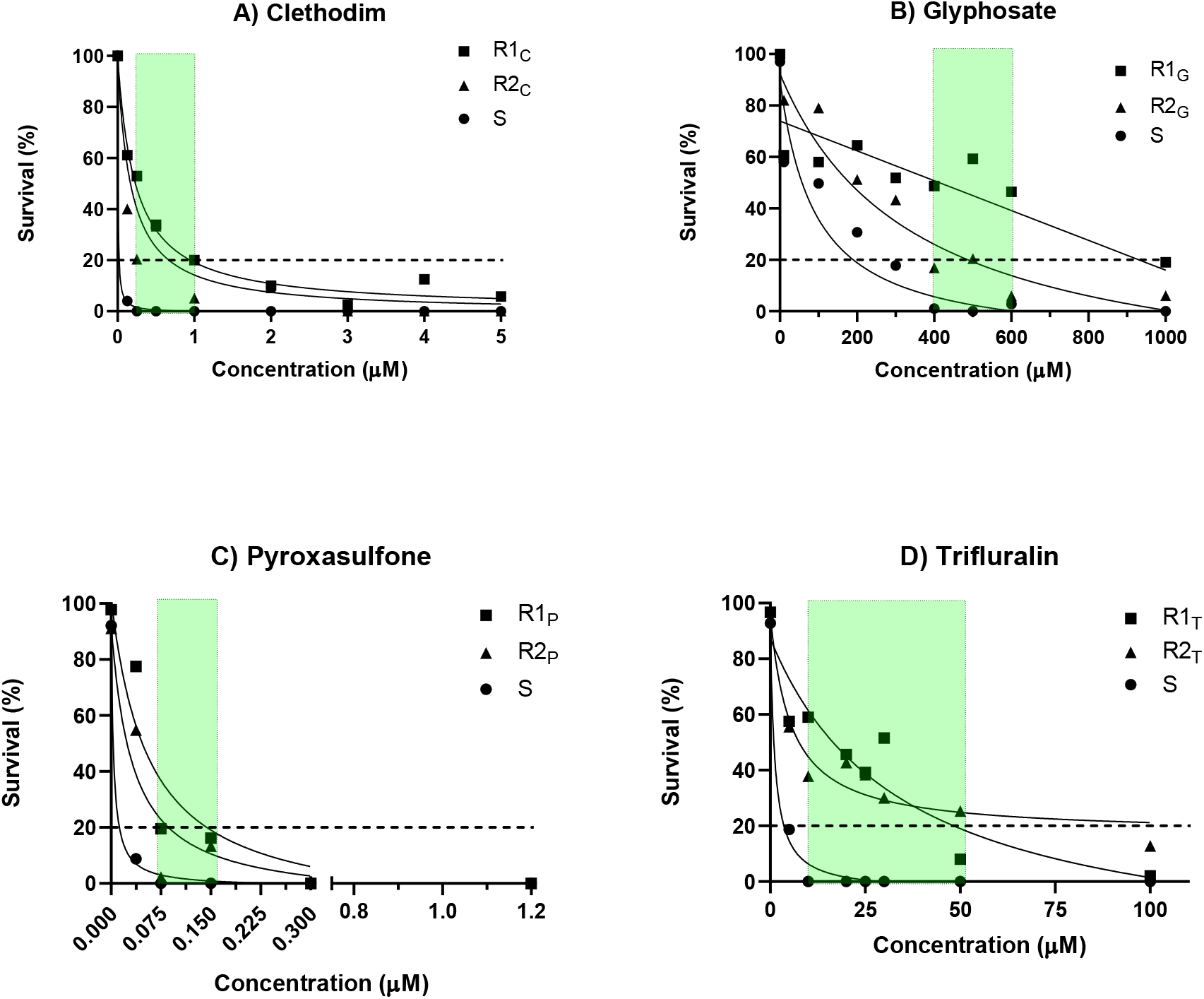
Dose-response curves for seeds of susceptible (S) and resistant (R) populations of *Lolium rigidum* germinated on agar containing various concentrations of (A) clethodim; (B) glyphosate; (C) pyroxasulfone or (D) trifluralin. Each point is the average of two independent experiments, each with three replicates. Continuous lines are survival response estimated with a three-parameter regression model (Y = d / 1 + exp b(log(x) − log(e)). The dotted line represents a discriminating threshold to identify a highly resistant population (> 20% survival). The transparent green area defines a range of herbicide concentrations that could be used to effectively and rapidly detect herbicide-resistant phenotypes (0% survival of S versus 2 – 60% survival in R).

**Figure 3.**
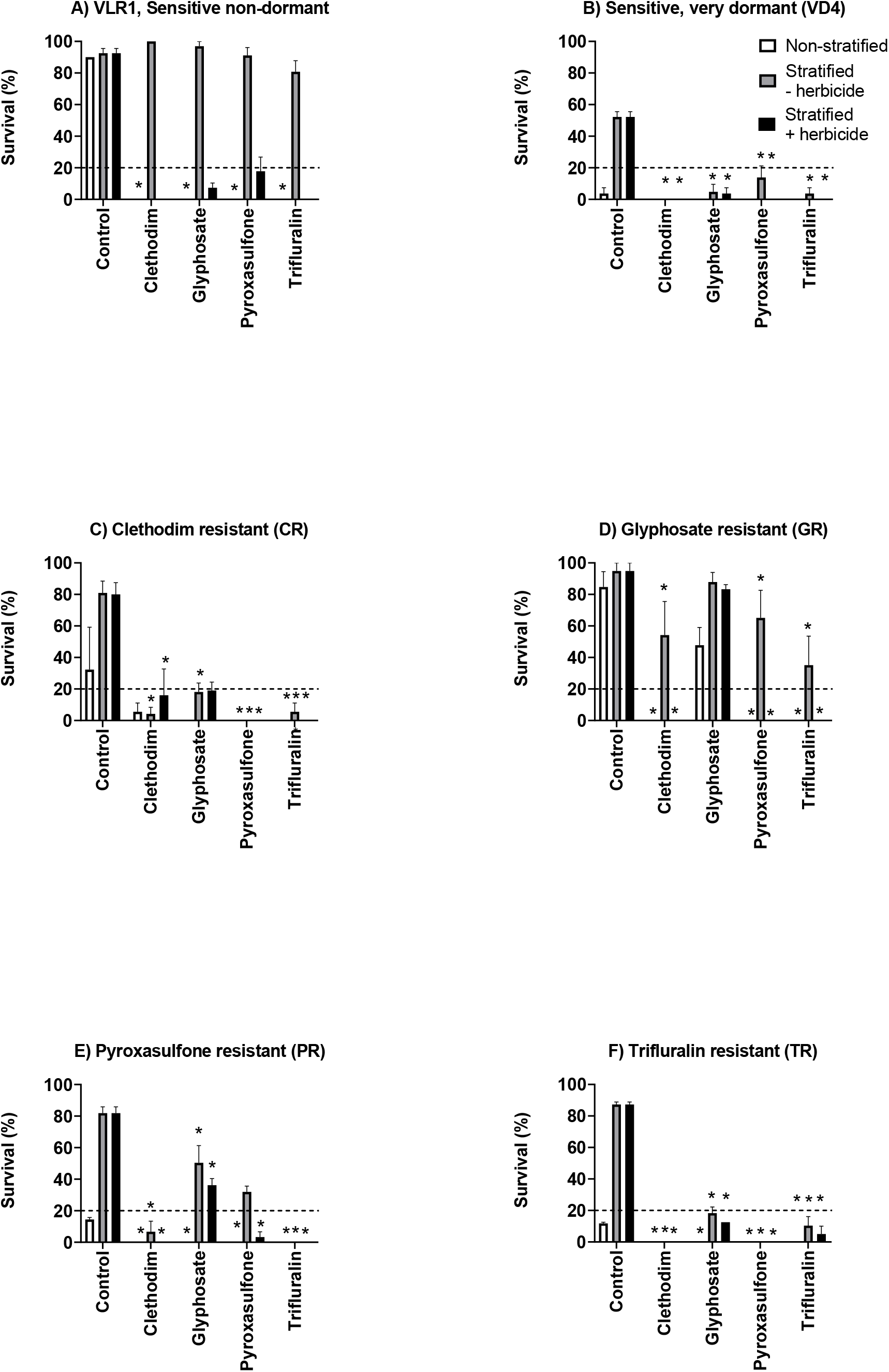
Apparent resistance levels (measured as the percentage of seeds germinating and producing a coleoptile of ≥4 cm) in A) non-dormant susceptible population S; B) dormant susceptible population VD4; C) clethodim-resistant population CR; D) glyphosate-resistant population GR; E) pyroxasulfone-resistant population PR; and F) trifluralin-resistant population TR germinated directly on herbicide-containing agar for 7 d (white bar), dark-stratified on plain agar for 21 d before being transferred to herbicide-agar for 7 d under germination conditions (grey bar); or dark-stratified in the presence of herbicide for 21 d with subsequent germination occurring on the same herbicide-containing media (black bar). Herbicide concentrations were 2 μM clethodim, 400 μM glyphosate, 0.150 μM pyroxasulfone or 20 μM trifluralin. Herbicide treatments resulting in significantly (*P*<0.05) different germination to the corresponding control are marked with an asterisk; the dotted line at 20% germination represents a high apparent resistance level. Values are means ± SE (n = 3).

### 3.2 Validation of the agar test with field populations

Agar and traditional pot experiments were conducted to compare the survival response of a large number (100) of field populations collected from the Western Australian wheat belt. As was expected from previous surveys (Busi and Beckie, 2020; Owen and Powles, 2018), the overall frequency of resistance to trifluralin and clethodim was >40% of populations at the discriminating dose in agar and at the field rate in pots. There were only five cases of resistance to glyphosate and no cases of resistance to pyroxasulfone; as a consequence of these low frequencies of resistant samples, only the data for clethodim and trifluralin showed significant correlations between percentages of survival in agar-based vs. pot-based resistance tests (Table 3). The agar test is thus able to discriminate characterized populations of annual ryegrass resistant to four different modes of action including clethodim (acetyl-CoA carboxylase inhibitor), glyphosate (5-enolpyruvylshikimate-3-phosphate synthase inhibitor), pyroxasulfone (very-long-chain fatty acid elongase inhibitor) and trifluralin (cell division inhibitor), and discriminate uncharacterized field populations to at least clethodim and trifluralin. However, specific calibration of herbicide concentrations will probably be required for each different weed species tested. In black grass (*Alopecurus myosuroides* Huds.), for example, it has been shown that false-negative results are observed with seed-based assays in which populations with resistance mutations at position 1781 of acetyl-CoA carboxylase, clearly resistant to field rates of herbicide in pot studies, remain susceptible when tested at the seed stage (Délye et al., 2008).

**Table 3.** Correlation coefficients (r) of herbicide survival response obtained in pot-based versus agar-based studies by screening 100 populations of annual ryegrass. ^ns^, not significant; **, *P* < 0.01; ***, *P* < 0.001.

|  |  | Agar |  |  |  |
| --- | --- | --- | --- | --- | --- |
|  |  | Clethodim | Glyphosate | Pyroxasulfone | Trifluralin |
| Pots | Clethodim | 0.426*** |  |  |  |
|  | Glyphosate | -0.034<sup>ns</sup> | 0.161<sup>ns</sup> |  |  |
|  | Pyroxasulfone | 0.017<sup>ns</sup> | -0.092<sup>ns</sup> | -0.098<sup>ns</sup> |  |
|  | Trifluralin | 0.076<sup>ns</sup> | -0.023<sup>ns</sup> | -0.211<sup>ns</sup> | 0.337** |

### 3.3 Interaction between seed dormancy and the agar test

In all experiments, the response of seeds germinated on the laboratory window sill were similar to those germinated in the growth cabinet (Supplementary Table S2), so only the latter results are shown. Several methods of chemical or mechanical scarification or chemical stimulation (Supplementary Table S1) were tested on seeds with primary or conditional dormancy to determine if the requirements for dark-stratification and alternating light and temperature could be overcome. In the absence of any treatments, germination of conditionally-dormant seeds after 7 d was 50%, whilst only 5% of the seeds with primary dormancy had germinated in this time (Supplementary Figure S1). No scarification or stimulation method was successful in increasing germination to >70% in the first 7 d (grand average: 28%) (Supplementary Figure S1).

The germination of highly dormant seeds placed directly under non-stratified germination conditions in the presence or absence of herbicide was too low after 7 d for the results to be reliable (Figure 3B, E, F). Dark-stratification of seeds in the absence of herbicide improved the germination percentage of the dormant populations once they were transferred to herbicide-agar under germination conditions, but resulted in premature germination of the less-dormant populations so that few ungerminated seeds were available for transfer (Figure 3A, D). All dormant populations stratified on herbicide-free agar and transferred to herbicide-containing agar for germination appeared susceptible to clethodim and trifluralin but resistant to glyphosate, with the exception of the VD4 population, which appeared susceptible to all herbicides (Figure 3B, C, E, F). Only the pyroxasulfone-resistant population appeared resistant to pyroxasulfone after stratification on herbicide-free agar (Fig. 3E).

Dark-stratification of seeds in the presence of herbicide resulted in apparent resistance to glyphosate in the clethodim-, glyphosate- and pyroxasulfone-resistant populations, and to pyroxasulfone in the glyphosate- and pyroxasulfone-resistant populations (Figure 3). No populations stratified on clethodim or trifluralin appeared resistant to these herbicides, even those populations previously characterized as resistant. Futher work would be needed to fully elucidate such a complex link between seed dark-stratificaiton and a putative induced or reduced ability to survive discriminating herbicide concentrations.

Therefore, the first constraint to the adoption of this agar test as a portable kit for farmers to use on-farm is seed dormancy. Seed that does not germinate due to dormancy could be interpreted as being herbicide-susceptible and give false negative results (Moss, 1995). In the current study, the characterized populations that were used to develop the methodology were fully after-ripened and had little or no dormancy (>96% germination within a week). None of the rapid dormancy-breaking treatments tested on dormant annual ryegrass seeds were able to short-circuit the requirement for months of dry after-ripening or weeks of dark stratification. Dark stratification itself had a tendency to confound the results, with several populations appearing to be glyphosate-resistant when previous pot tests had shown that they were not. The requirement for low-dormancy seeds to ensure accurate results means that freshly-collected seeds need to have their dormancy level assessed and then be left to after-ripen if necessary, which erodes the time advantage of the agar test compared to pot experiments.

### 3.4 Stability of herbicides in stock solutions and agar plates

There was no degradation of the 10 mM pyroxasulfone and glyphosate stocks stored at 4°C across the 236 days of the experiment, but some slight (~15%) loss of trifluralin and clethodim was observed (Supplementary Table S4). When diluted in agar, clethodim remained stable when stored in the dark at room temperature, but 50 – 70% was lost between 5 and 10 d in the other two treatments (Figure 4). Glyphosate remained stable when stored in the dark at either room temperature or 4°C, but the concentration in plates stored under ambient light and temperature decreased by around 10% over the 10 d (Figure 4). There was no significant change in pyroxasulfone concentration over time under any conditions. In contrast, there was an almost complete loss of trifluralin in the first 5 d in plates stored at room temperature in the light or dark, but a much smaller decrease (around 25%) in the samples stored at 4°C (Figure 4).

**Figure 4.**
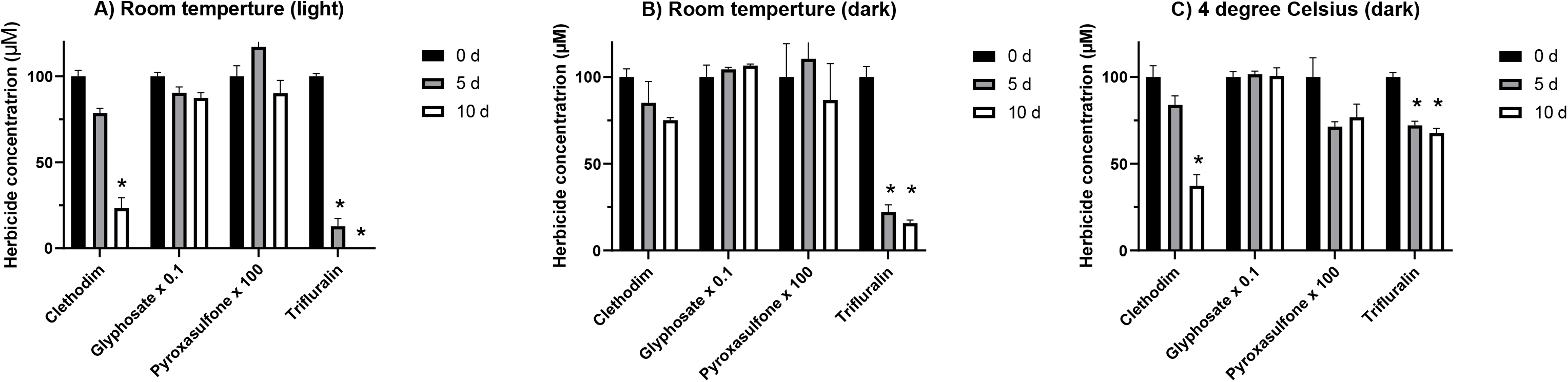
Stability of herbicides in agar. Herbicides incorporated in agar at concentrations of 2 μM clethodim, 400 μM glyphosate, 150 nM pyroxasulfone or 20 μM trifluralin were stored at (A) room temperature under ambient light; (B) room temperature in the dark; or (C) 4°C in the dark and monitored at 0, 5 and 10 d after the start of the experiment. Values are means ± SE (n = 3); asterisks above bars denote herbicide concentrations that are significantly (*P* < 0.05) different to the corresponding starting concentration (0 d).

This demonstrates that the other major limitation of the agar test is the rapid loss of trifluralin, and clethodim to a lesser extent, upon dilution in agar. With the necessity of using trifluralin-containing agar almost immediately and clethodim-containing agar within 5 d of preparation, the shelf-life of an agar test kit would not be sufficient for the kit to be sent out and used by farmers. Interestingly, trifluralin is able to prevent growth of trifluralin-susceptible seedlings after 42 d (Supplementary Table S2), even when trifluralin in the bulk agar is depleted after 5 d (Figure 4). This result could be explained by binding of trifluralin to the seeds, maintaining a high local concentration even when the herbicide evaporates from the surrounding agar. Dinitroaniline herbicides are known to bind strongly to organic matter (Carringer et al., 1975; Peter and Weber, 1985), particularly lipids, and therefore are able to accumulate in the cell membrane (Brand and Mueller, 2002).

### 3.5 Uses of the agar test

The classical laboratory-based agar test is readily transferable to other weed species and herbicides with other modes of action (Kaundun et al., 2011), and has often been used in research on herbicide resistance mechanisms. For example, it has been shown that an agar test can discriminate between target and non-target site resistance to diclofop-methyl by comparing shoot vs. root elongation (Preston and Powles, 1998). Similar methods have been used to screen resistance phenotypes in annual ryegrass populations and subsequently identify the resistance-conferring genes to acetolactate synthase, acetyl-CoA carboxylase and mitosis-inhibitor herbicides (Burnet et al., 1994; Chen et al., 2018a; Kaundun, 2014; Powles et al., 1998; Tardif et al., 1993; Tardif and Powles, 1994; Yu et al., 2007b). Although unsuitable for on-farm resistance testing, the agar test can be used as a complement to large-scale pot-based resistance screening. For example, it can confirm resistance in selected populations in a dose-response experiment with a greater range of herbicide concentrations. In addition, an agar test can be used to compare phenotypes of strongly- vs. weakly-resistant populations, and is particularly suited to the study of pre-emergence herbicides as individual seeds are visible and they can be fully monitored.

## 4 Conclusion

In conclusion, the agar test can be considered as a resistance detection method that is complementary to traditional pot-based tests, can be developed for any herbicide and is especially useful for detailed studies of populations identified as resistant. However, the agar test described herein is not suitable for portable use on-farm due to the complicating factors of seed dormancy and herbicide degradation. Greater adoption of pot-based resistance testing services would help end-users to deploy the most effective herbicide treatments by alternation of different mode of action which in turn would help ensure the long-term preservation of weed control resources.

## Acknowledgements

This work was funded by the Grains Research and Development Corporation (GRDC) as part of the Innovation Program 2018 - Accelerating Innovative Technologies (managed by Manjusha Thorpe).

